# Bee communities in canola are affected by landscape context and farm management

**DOI:** 10.1101/2021.11.29.470453

**Authors:** Rachel L. Olsson, Vera W. Pfeiffer, Benjamin W. Lee, David W. Crowder

## Abstract

Bees are key pollinators that promote greater yield and seed quality of oilseed crops such as canola. Canola acreage has increased over 1,000% in the past decade in the Pacific Northwest USA, providing a major pulse of sugar-rich nectar and pollen resources that may affect bee health and community structure. However, because canola does not require insect pollination for seed production, few studies have examined the biodiversity of pollinators taking advantage of these resources, or the floral traits of canola that affect pollinators across variable landscapes. Here, we conducted pollinator surveys at canola farms across the inland Northwest region of the USA. We surveyed bee biodiversity and abundance, and assessed how these metrics correlated with landscape context, canola production practices, and floral traits of various canola varieties. We found that bee communities differed between sites and across growing seasons, with sweat bees more abundant later in the season, and mining bees more abundant earlier in the season. We also found that bees were more abundant overall on farms with less floral nectar and with less developed landscape surrounding the sampling area. Bee diversity was greater in spring canola than winter canola, and floral traits were also correlated with differing bee community diversity. This research provides important information for canola growers and land managers and offers a framework for future research in pollinator management in the inland Northwest.

## Introduction

Bees contribute pollination services to the vast majority of food, fuel, and oil crops consumed worldwide (Klein et al. 2007). There are an estimated 25,000 species of bees around the world, most of which are solitary (Michener 2007). Bees feed on nectar as adults and collect pollen to feed to their developing young. Many bees are generalist foragers and feed on a wide array of different flower types, while others can complete development on single species of plants (Cane et al. 2006; Jha et al. 2013). However, shifts of natural land into agriculture, urban spaces, and other human-mediated landscape types have led to dramatic declines in the availability of floral resources across the globe, which has resulted in bee population declines (Goulson et al. 2015).

The Palouse region of the USA, which comprises eastern Washington and northern Idaho, has seen extensive agricultural intensification over the past century (Black et al. 1998). The Palouse is dominated by dryland cereal and legume production and is among the most productive agricultural regions in the world. However, the Palouse was historically a prairie landscape with diverse flowering plants that provided food resources for many wild bee species. Indeed, over 170 species of wild bees have been found in surveys of remnant Palouse prairie habitats, despite the fragmented nature of these patches (Rhoades et al. 2017). Yet, due to the homogenization and simplification of the Palouse, food resources for bees are sparse. Wheat does not require insect pollination and does not produce insect-attractive resources such as nectar. As the acreage of wheat has increased over the past century, and prairie remnants have become more scarce and fragmented, the food and refuge area available for bees has declined (Black et al. 1998).

Over the past decade, farmers in the Palouse have increasingly grown canola to diversity crop rotations (Pan et al. 2016). The acreage of canola in eastern Washington, for example, has increased from 5,000 acres in 2007 to over 70,000 acres in 2019 (WOCS 2017, USDA-NASS 2019). Canola does not require insect pollination, but studies show that seed yield can increase by up to 40% when insect pollinators are present (Morandin and Winston 2005, Bommarco et al. 2012). Canola can be grown as a fall-planted or as a spring-planted crop, depending on climate (Koenig et al. 2011, Esser and Hennings 2012, Pan et al. 2016). Both winter and spring canola flower for several weeks, providing a large pulse of floral resources for bees (Bjerke et al. 2019). Canola bloom also typically coincides with emergence of bumble bee queens and early season solitary bees. If both winter and spring canola are grown in proximity, canola nectar and pollen are available throughout bee’s entire life cycle. For social bees, the high concentration of floral resources provided by canola can boost colony growth rates, and for solitary bees canola may provide all the necessary resources to support entire broods. However, in the Palouse it remains relatively unknown which bees visit canola, or which canola traits attract bees.

Here, we conducted an study across Palouse landscapes to identify which bee groups use canola as a food resource, and which traits and varieties of canola attract particular bee groups. Studies show that bees have differences in preference for flowers based on different floral traits, including flower size and nectar volume or quality (Finke 2012, Ibanez 2012, Carruthers et al. 2017). As farmers use variable practices to produce canola, such as irrigation and tillage, we also assessed how management practices affected bee abundance and diversity. As canola is grown throughout variable landscapes of the Palouse, we further investigated how the context of landscapes surrounding canola fields correlated with bee abundance and diversity. We predicted that pollinator communities would vary based on land use type, farm management, and plant traits such as flower size and nectar quality. Our study provides insight into factors affecting bee communities in canola, and can be used to guide pollinator conservation and agronomic tactics.

## Materials and Methods

### Study system

Canola is a flowering member of the Brassicaceae family that is grown to produce seeds used for food, feed, and fuel. Canola is an industry trade name that applies to *Brassica rapa, B. napus*, or *B. juncea*, all of which have a low proportion of erucic oil (less than 2%) in the seed (Canada 2017). In the Palouse region, the climate is such that both winter and spring varieties may be grown successfully. Winter canola is planted in early fall, allowed to grow, and is winter-stunted by cold temperatures. Once spring temperatures rise, the plants resume growth and begin flowering in mid-spring before harvest in mid-summer (Esser and Hennings 2012). Spring canola crops are planted once the soil has thawed, typically in early spring. These plants will begin flowering in early summer and then harvested in late summer (Pan et al. 2016).

The choice of growing winter or spring canola depends on factors such as soil moisture, seasonal temperatures, and crop rotations. Moisture limited sites will most frequently grow winter canola because most precipitation falls in autumn when plants are establishing. Growers who have access to irrigation might plant either winter or spring canola, and those who live in higher rainfall areas often grow spring canola. However, whether winter or spring canola have variable effects on bee populations, or whether varieties of each differ in their attractiveness to different bees, is unknown. We predicted that different communities of bees would visit winter season canola, which blooms from late April to late May, than spring canola, which blooms later (mid-June to early July) (Olsson et al. 2021). We also expected to see larger populations of bees fields of canola with higher nectar volume and sugar, and visiting plants with larger, more attractive flowers (Carruthers et al. 2017, O’Brien and Arathi 2018, Adamidis et al. 2019).

Farmers in the Palouse also use variable production practices to grow canola. Many farms in this region practice no-till or low-till farming practices, but these techniques require heavy equipment to drive over soil, which can cause compaction and disturbance. These disturbances could have an impact on the nesting suitability for the soil, or if tilled, could entirely destroy wild bee nests (Ullmann et al. 2016). Some canola growers also use irrigation to supplement natural rainfall, while many do not. The effects of these agronomic practices on bee populations, and the attractiveness of canola flowers, might differ across variable landscapes of the Palouse. For example, in areas with more natural habitat, there may be larger pools of bee species available to respond to canola (Kennedy et al. 2013; Lichtenberg et al. 2017). However, effects of agronomic practices and landscape context on bee communities in canola are unknown.

### Collecting bees

To address our study questions, we sampled the community of pollinators at canola farms distributed across eastern Washington, northern Idaho, and northeastern Oregon in 2018 and 2019 (Fig. 1). Since producers grow canola in rotations, not all farms visited were the same each year, though several farms were revisited. We visited farms during the peak flowering period, as determined by participating growers. Peak flowering occurred when the most flowers were blooming at one time, and typically occurred five days after flowers began blooming.

**Figure 1.**
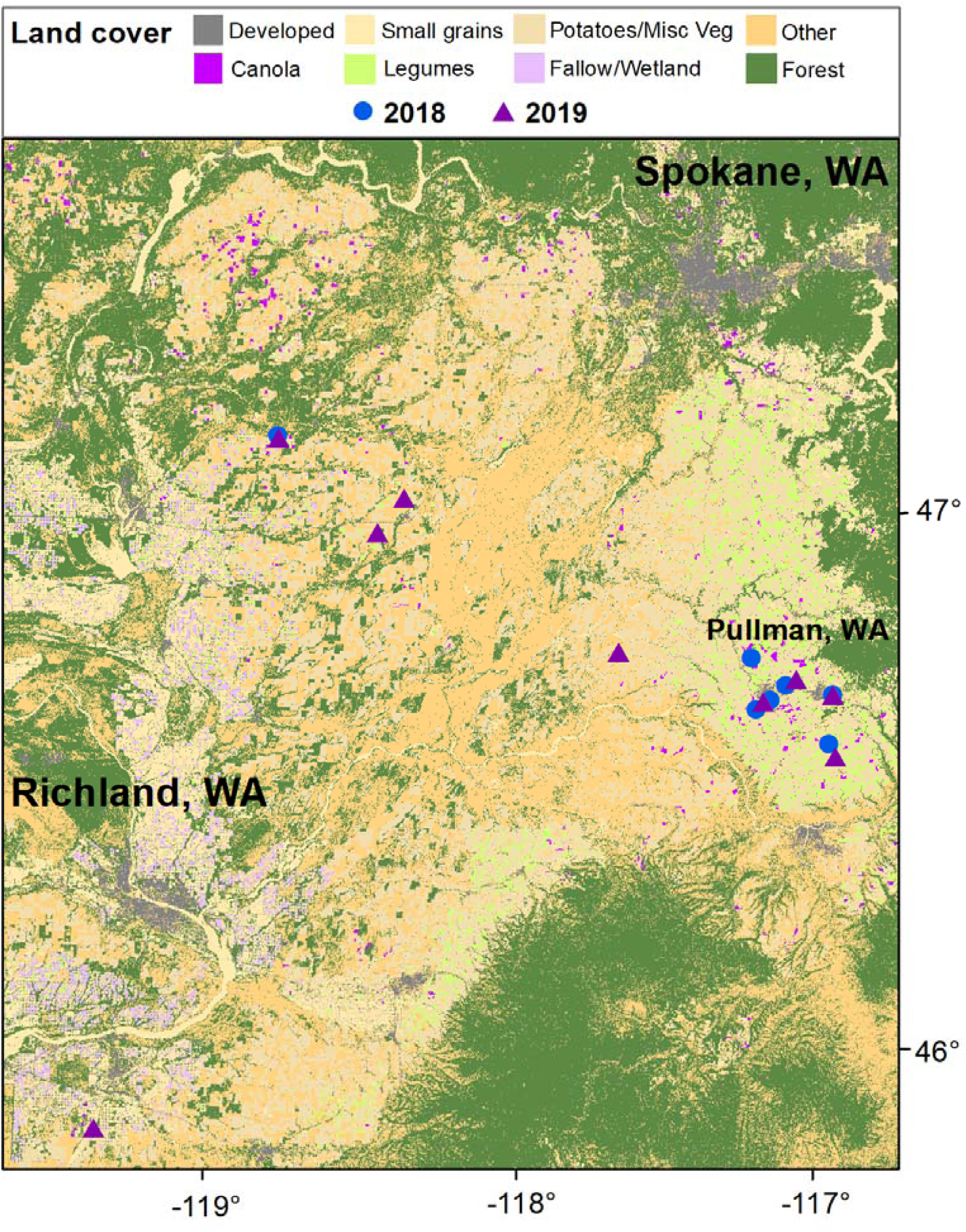
Map showing farm locations in 2018 and 2019 with landscape types across the range of our study system (Oregon, Washington, Idaho)

We collected bees using three methods: (i) blue vane traps, (ii) bee bowls, and (iii) aerial netting. Blue vane traps were placed about 10 m apart along a transect bordering one of the edges of the canola field. Blue vane traps were suspended by twine from a bamboo tripod, so the trap hung at the level of canola flowers. Bee bowls were placed between blue vane traps 2.5 m apart. One bowl of each color was randomly selected (blue, yellow, white) and placed between the blue vane traps, for a total of 6 bowls (two of each color). Bowls and blue vane traps contained soapy water and were placed at field edges between 08:30 and 09:00. Traps were removed 6 h later. All contents of traps were strained through a fine mesh filter, rinsed with clean water, and insects were placed in bags filled with 70% ethanol. Bee bowls and blue vane traps are known to attract most types of bees (Rhoades et al. 2017). However, we also included two 15 min bouts of aerial netting at each site to more fully sample the bee community. We made one netting trip in the morning and one in the afternoon. For each netting bout, we walked along the field edge and between rows, netting every bee seen on a reproductive part of a flower. Bees were moved from the net into a 50 mL vial filled with 70% ethanol for storage until later identification.

To complement our passive and active sampling methods, we made visual observations of bees visiting flowers. This allowed us to determine which bees were visiting canola flowers, and not just bees in the vicinity of canola fields. For this, we walked between field rows, stopping every 5 m to cast a 1 × 1 m visual square over a patch of canola flowers. We then spent 30 sec noting every bee that visited the reproductive part of a flower within that square. On-the-wing identification to species was infeasible, so we visually identified bees to morphotype. These morphotypes included mining bees (Andrenidae), bumble bees (Apidae: Bombus), carpenter bees (Apidae: Xylocopa), green sweat bees (Halictidae), honey bees (Apidae: *Apis melifera* L.), longhorned bees (Apidae: Melissodes), masked bees (Colletidae), megachilid bees (excluding blue orchard bee; Megachilidae), blue orchard bees (Megachilidae: Osmia), medium to large sweat bees (Halictidae), and small sweat bees (excluding green sweat bees; Halictidae).

### Processing and identification

Once returned to the lab, all bees were rinsed in ethanol, dried, and then pinned for later identification. Bees were sorted based on data and sampling location, as well as by method of collection. Bees were then identified to morphotype using the same broad groupings that had been used during the visual field observations.

### Canola petal measurements

During our field sampling, we took samples of canola petals to determine the average area of the petals. For each of eight plants, haphazardly selected in the field, we took two petals from each of three flowers, totaling six petals per plant, and 48 petals per field. We taped these petals to a data sheet that included a size standard black box printed on it. This box measured 46 × 31 mm, or 1,426 mm2. We scanned the data sheets and used ImageJ software (Rasband, 2018) to measure the size of each individual petal in pixels, and used the size standard to calculate the area in mm2 of each petal (Glozier 2008).

### Canola nectar measurements

While in the field, we also took nectar measurements from flowers of six randomly selected plants. Using a 10 μL microcapillary tube, we extracted nectar from six flowers on a single plant, drawing nectar from each floral nectary. We measured the length of the nectar in the tube using digital calipers as a proxy for volume, due to the standardized volume of the tube. Then we added 4μL of deionized water to the nectar sample and assessed nectar sugar concentration using a refractometer (dilution allowed us to assess all samples) (Farkas et al. 2012). Diluted values were then converted to actual nectar sugar concentrations.

### Data analysis

All data were analyzed using the R Statistical Programming software (R Core Team, 2019). Figures were generated using the ‘ggplot2’ and ‘ggmap’ packages (Kahle and Wickham 2013, Wickham 2016) and the ‘viridis’ color package (Garnier 2018). Community ecology analyses were performed using the ‘vegan’ package (Oksanen et al. 2019). To gather landscape data, we mapped and extracted croplands data by mask in ArcGIS10.7 using 2 km buffers around our study sites, and used the zonal metrics package to calculate class area, percentage, and number of patches (Mohler et al. 2009, USGS 2018, 2019, Fig. 1).

For analyses, we first used linear mixed effects models to examine effects of canola type (winter or spring) and canola variety (nested within canola type) on petal size, nectar volume, and nectar sugar concentration. These variables were nested because particular varieties were associated with only winter or spring canola. In each of these analyses, farm site was included as a random effect. At many of our sites, we were unable to collect a measurable volume of nectar. For these sites, nectar volume was determined to be 0, but these sites were excluded from the analysis of nectar volume and nectar sugar concentration. Second, we used a generalized linear mixed model with a Poisson transformation to measure effects of tillage, irrigation, canola type (winter or spring), canola variety (nested within type), canola petal size, nectar volume, and area of three major land cover types (canola, developed, and legumes) on bee abundance (Chambó et al. 2017). Farm was included as a random effect in these models.

Finally, as bee communities may respond to landscape context, agronomic practices, and canola traits through direct and indirect pathways, we used structural equation models to test several *a priori* predictions (Fig. 2). First, we assumed that canola type (winter or summer), tillage practices (conventional, conservation, no-till) and irrigation (present or absent) could directly affect nectar sugar content, nectar volume, and petal size (Pan et al. 2016, Adamidis et al. 2019). Second, we assumed that canola type, tillage, irrigation, and canola traits (nectar sugar content, nectar volume, petal size) could affect bee abundance and biodiversity (measured as Shannon’s index) (Ullmann et al. 2016). Third, we assumed that landscape context could affect bee abundance and biodiversity (Torne-Noguera et al. 2014). Finally, we assumed that nectar sugar content and nectar volume were correlated traits, where we could not assume a particular direction of the effect (Fig. 2) (Wiemer et al. 2012). Within R, we built this a priori model using a series of linear regression models, and then used the piecewiseSEM‘’ package to construct the final path model (Lefcheck 2016). For analysis, nonsignificant paths that reduced AIC were dropped, and paths were added if models without them were rejected via directed separation tests and the relationships were biologically feasible (Lefcheck 2016).

**Figure 2.**
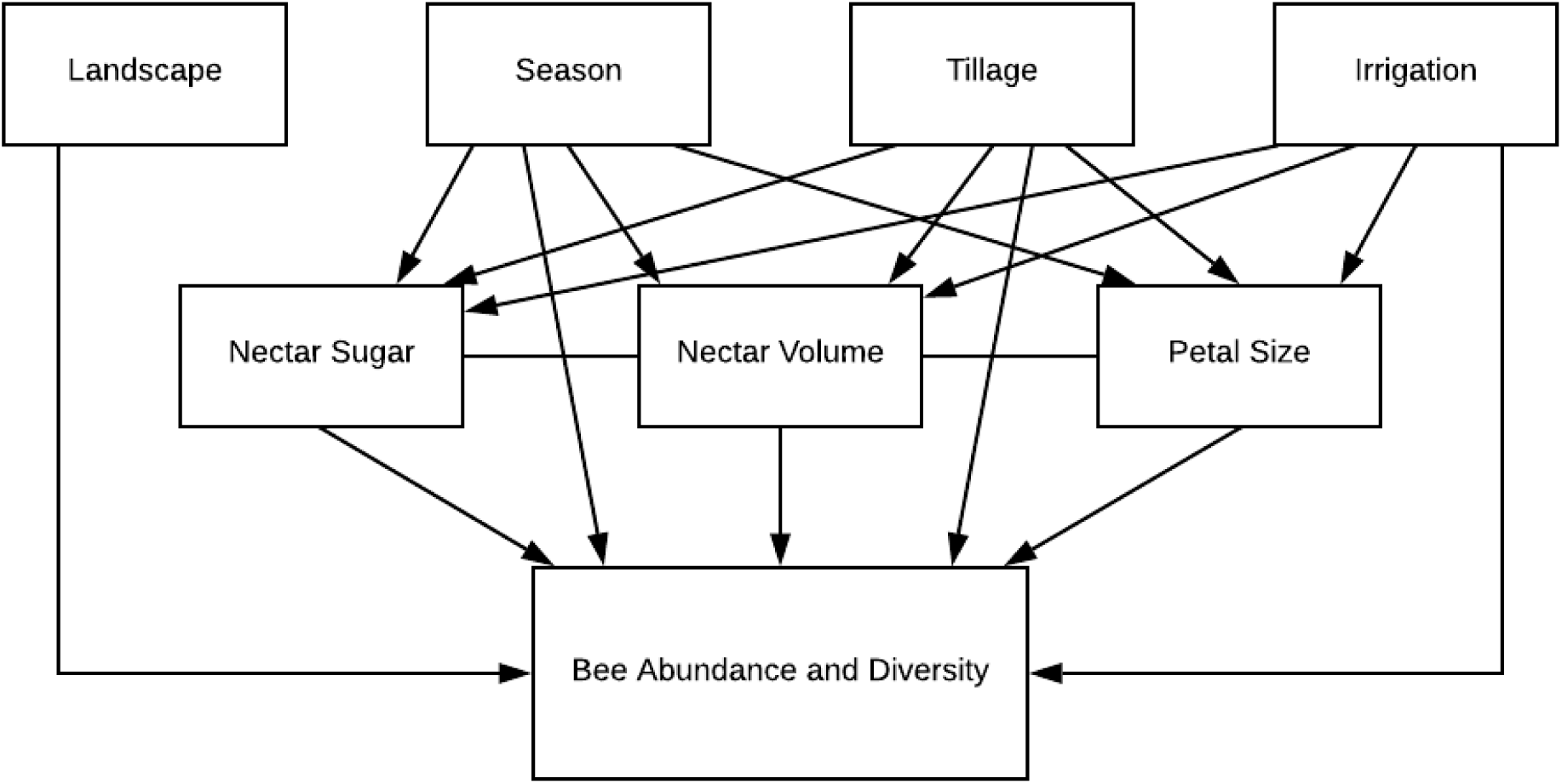
*A priori* structural equation model showing hypothesized predictor variables (top, middle), and hypothesized response variables (middle, bottom)

## Results

We measured plant traits and sampled the bee community at 13 sites over two years. Across the two years of sampling we collected a total of 3,287 bees across five families and 13 genera (Fig. 3A). The majority of these bees were collected at a single farm in 2018; at this site we collected over 1,700 individuals (site Du18w, Fig. 3A). We recognized the possibility that this strong outlier might exert high leverage on the overall results, so during analysis, we analyzed all data with and without that site. However, we observed the same trends when the data were excluded, so the outlier site was included in all further analyses.

**Figure 3.**
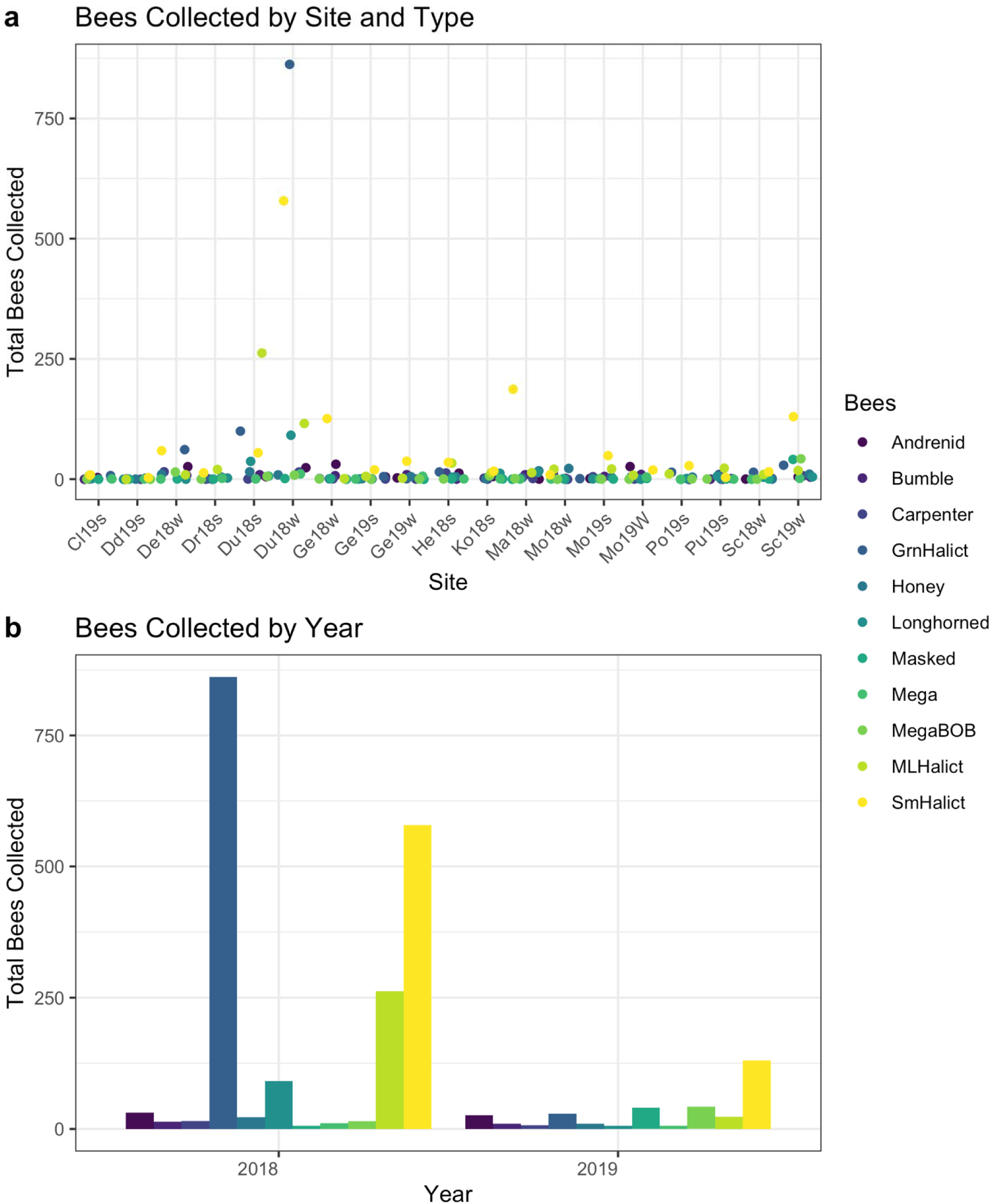
(A) Bees collected during each sampling event over 2018 and 2019 and (B) bee abundance measurements collected by year and sorted by bee morphotype.

### Nectar, petal size, and bee guilds

Canola petal sized varied significantly across the different varieties (Fig. 4A). The Edimax variety had the largest petals (98.4 ± 1.99 mm2), followed by varieties 05.WC.34.5 (74.8 ± 1.99 mm2) and InVigorL233P (75.3 ± 1.99 mm2). These varieties had petals that were significantly larger than the most common variety, HyClass930 (70.6 ± 0.94 mm2), and the Winfield-Hyclass variety (72.4 ± 1.77 mm2). The variety with the smallest petal size was 46W94 (65.6 ± 1.99) (Fig. 4A). Nectar volume was another factor that varied across the different sites and across varieties, although several sites did not have measurable nectar (Fig. 6). As with nectar volume, because we were unable to collect a measurable amount of nectar, we were unable to measure the sugar concentration at many of our sites (Fig. 7).

**Figure 4.**
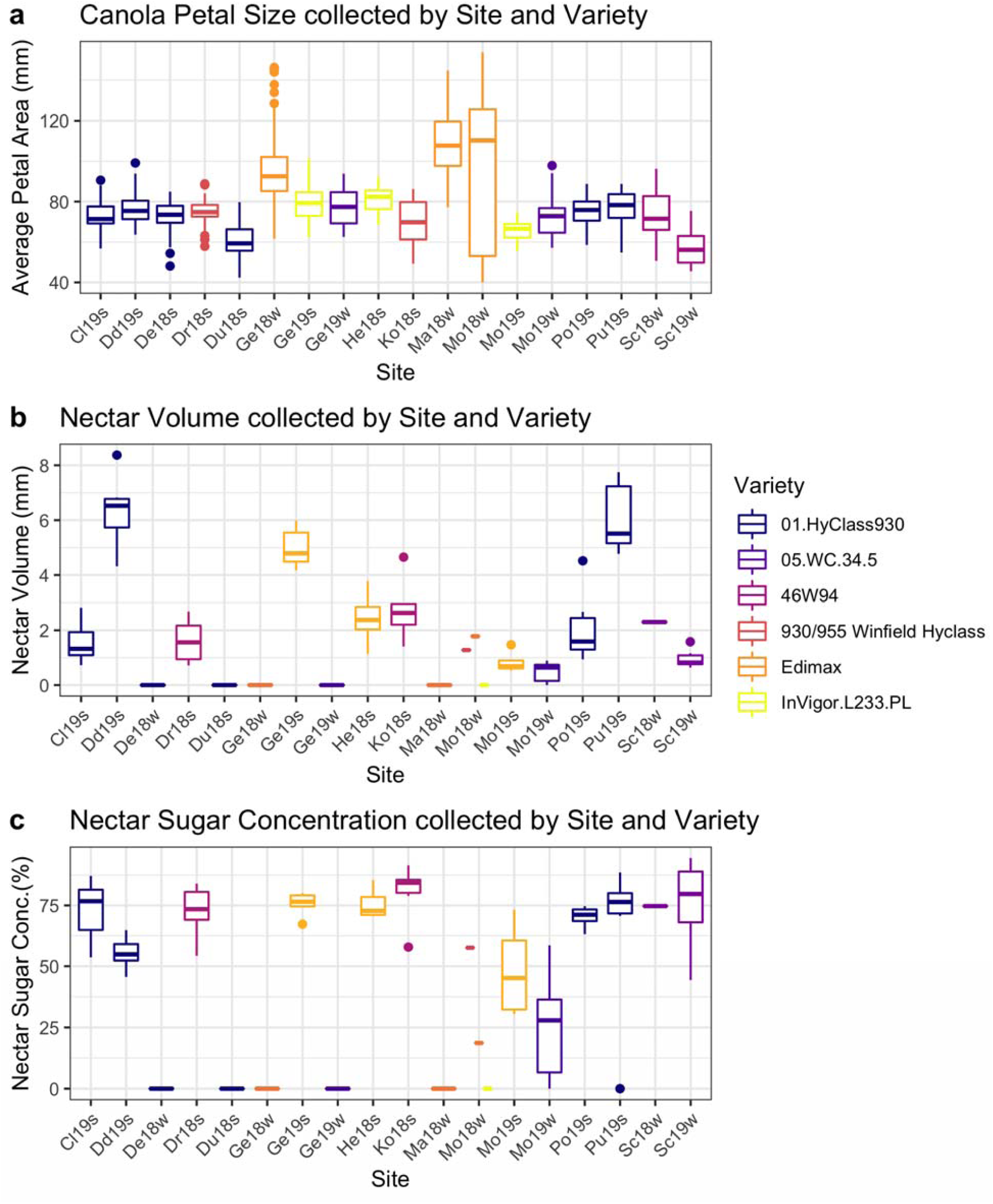
(A) Canola petal variety and average size collected at each farm visit in 2018 and 2019. (B) Canola nectar volume (mm) by variety and farm in 2018 and 2019. (C) Canola nectar sugar concentration (%) by variety and farm in 2018 and 2019

Across all our sites, pooled across both years, we found that the most abundant bee groups were green sweat bees and small sweat bees (both Halictidae) (Fig. 4). Small halictid bees were the most abundant group (t198 = 2.72, P = 0.0071) across all farms, followed by green bees (t198 = 2.16, P = 0.032). All other bee groups were not significantly different in abundance across farms. The land coverage 2km around farms did not affect bee abundance (F4,13 = 0.74, *P* = 0.56). Overall, bees were more abundant at farms with smaller petals and lower nectar volumes (generalized linear mixed model, Z = -9.52, P < 0.001 and Z = -2.01, P = 0.040, respectively).

### Environmental factors and bee abundance

There were significantly fewer bees in locations that used minimal tillage and no tillage, compared with conventional tillage (GLMM, Z = -7.34, P < 0.001 and Z = -5.25, P < 0.001, respectively). Bees were more abundant in irrigated than in non-irrigated sites (GLMM, Z = 5.81, P < 0.001). This was surprising because the irrigated fields were planted in winter canola, and bees had overall lower abundance in winter than in spring, though that result was not significant (GLM, Z = -0.02, *P* = 0.99). Bees were less abundant in areas with more developed land around the farm area sampled (GLMM, Z = -14.1, P < 0.001).

Our structural equation model corroborated many of the the trends seen in the simpler linear models (Fisher’s C = 3.68, *P* = 1.0, df = 14, Fig. 5, Table 1), but identified fewer factors driving significant relationships between environmental factors and floral and bee responses. Spring canola positively affected nectar volume (βstd = 0.68, P = 0.005), and increasing petal size negatively affected bee diversity (βstd = -0.56, P = 0.05). Increases in developed land around farms negatively correlated with nectar volume (βstd = -0.29, P = 0.17), and bee abundance (βstd = -0.69, P = 0.003, Fig. 5, Table 1).

**Figure 5.**
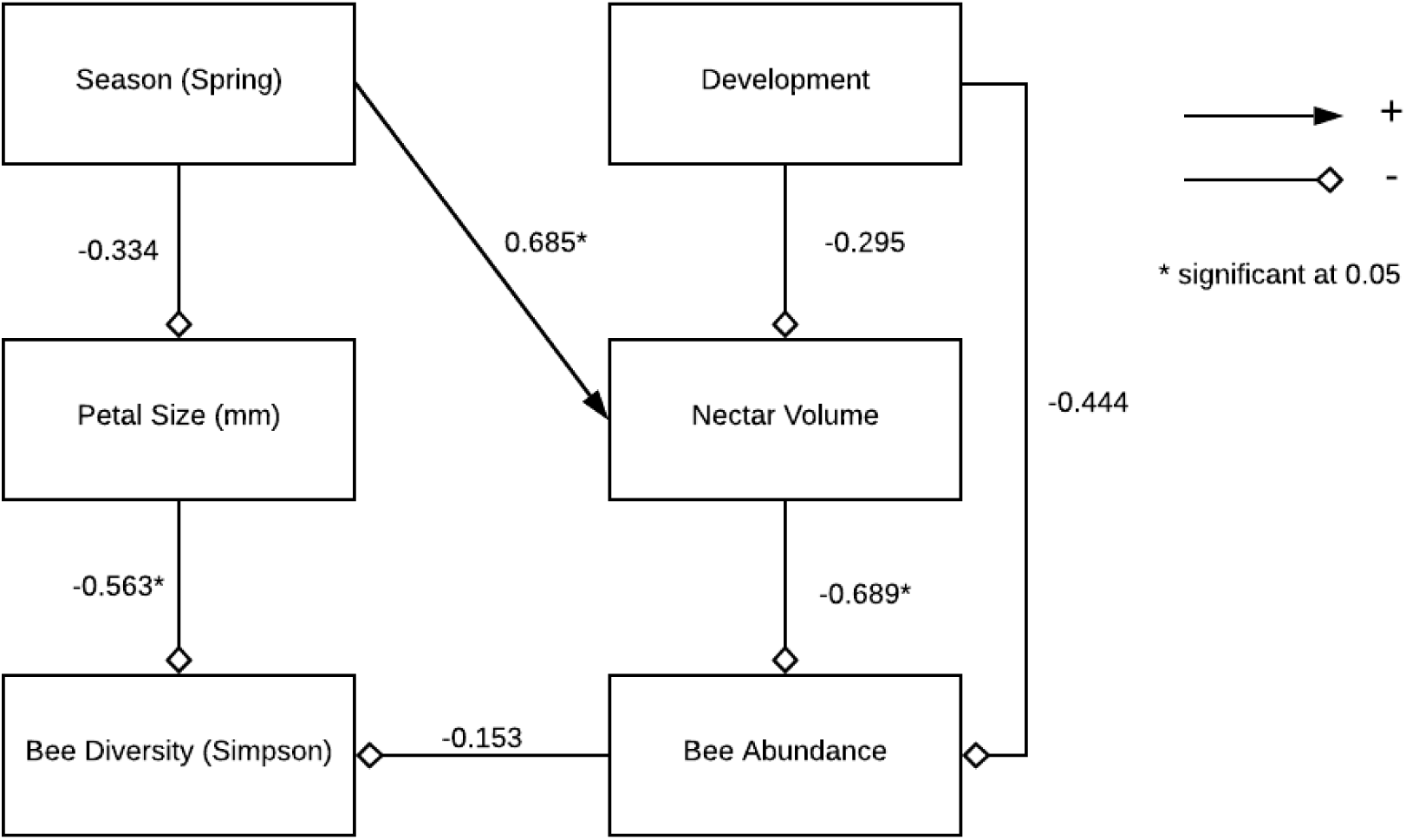
Supported model from confirmatory path analysis (Fisher’s C = 3.68, P = 1.0. df = 14).

**Table 1.** Standardized path coefficients and significance tests from accepted path model. Standardized coefficients (βstd) indicate relative magnitude and direction of effect.

| Path Coefficients |  |  |  |  |
| --- | --- | --- | --- | --- |
| Response | Predictor | $\beta_{std}$ | DF | P-value |
| Nectar Volume | Spring Variety | 0.68 | 14 | 0.0047 |
|  | Prop. Developed Landscape | -0.30 | 14 | 0.17 |
| Petal Size (mm) | Spring Variety | -0.33 | 13 | 0.22 |
| Simpson Diversity | Bee Abundance | -0.15 | 12 | 0.55 |
|  | Petal Size (mm) | -0.56 | 12 | 0.05 |
| Bee Abundance | Prop. Developed Landscape | -0.44 | 14 | 0.04 |
|  | Nectar Volume | -0.69 | 14 | 0.0029 |

## Discussion

Bees we collected represented many taxa, suggesting canola broadly supports pollinator communities. We were surprised, however, to see that bees were observed in higher abundance when nectar volume was lower. However, we believe that the overall average nectar volume was brought down by the number of sites where we could not collect measurable nectar. Although we were unable to collect this nectar, bees are much more adept at exploiting floral resources than humans, so we expect they may have taken advantage of these resources regardless of our ability to detect them. The high abundances of bees at fields with low nectar values could indicate that we arrived later in the day than the bees and were thus unable to collect nectar.

Although we did not discretely measure bee body size, the groups of bees that were seen most abundantly (green sweat bees, small sweat bees) are all smaller bodied bees. Although some studies show that larger flowers are more attractive to bees (Murcia 1990), our results align with other studies that show bees choose flowers that are appropriate for their body size. Body size is another factor in foraging distance, where larger bees are able to forage further from their nests, while smaller bees remain closer to gather resources (Gathmann and Tscharntke 2002). We collected many small bees, which would indicate they are foraging near their nests, located in or very close to the canola field we sampled. These sweat bees nest in the ground on an annual cycle, collecting provisions over the summer and overwintering in the nest (Goulet and Huber 1993). We did not expect to see increased bee abundances in farms with conventional tillage, as this is the most disruptive type of soil disturbance that could destroy bee nests (Ullmann et al. 2016). However, bees could have established new nests in the canola fields, and the disruption would not occur until the end of the season or the following spring during ground preparation for the next crop. Bees could have also been nesting in nearby fallow fields, field margins, or farm roads, which might act as habitat refugia in these highly disturbed areas (Greenleaf et al. 2007).

Canola has been bred for traits like larger seeds, more oil, and high protein content (Brown and Davis 2020). Canola is commonly a self-fertile crop, so little emphasis has been put into breeding for pollinator attractive trait like flower petal size or nectar content. Our study shows that bee diversity is promoted with increased petal size, which could lead to greater pollination (Halinski et al. 2018). If bees are attracted to flowers based on body size or other trait, a survey of wild pollinators might inform growers of a variety of canola with appropriately sized flowers to best take advantage of the wild pollinators nearby. We saw low abundances of bumble bees and other large-bodied bees on farms with relatively small petals, even though we witnessed bee activity during the time these flowers were blooming. It is possible that small flowers are not suitable for large bees, and thus are more able to be pollinated by a smaller bee. Alternately, if a grower is planning to stock honey bees near their canola field, they may need larger flowers to support a more robust bee like a honey bee. It is also important to note how much nectar is being produced, because if the amount is not suitable for large honey bee colonies, the bees could end up being depleted resulting in bee or honey losses to the beekeeper (Goodrich 2019).

We expected to see strong positive correlation between the land surrounding the farm site planted in canola or legumes and bee abundance and diversity. Neither of these were predictors of bee abundance, however, but the amount of land surrounding the site that was developed was a strong negative predictor of bee abundance and diversity. Developed land included buildings, roads, and other human-made structures or impenetrable surfaces. These habitat modifications reduce the suitability for bees by depleting floral resources and nesting sites, so it makes sense that this type of landscape would correlate with the observed abundance of bees (Goulson et al. 2015). Although canola acreage has increased in the Palouse, it still makes up a small proportion of the overall landscape (Fig. 1). We expect that due to the patchy nature of this crop, without suitable corridors to travel between canola fields, populations of most bees will be limited to the field nearest to their nesting site (Greenleaf et al. 2007). It has been previously demonstrated that it takes up to three years to see changes in wild pollinator communities as a response to plant landscape change (Petanidou et al. 2008). This understanding might provide support to growers looking to understand their pollination needs and resources, or when it might be useful to seek the services of a beekeeper rather than relying solely on wild pollinators.

Our goal in this study was to understand which pollinators are using canola as a floral resource, and what factors might be influencing the pollinator communities at individual sites Our research provides data to inform canola growers on potential management practices that could encourage pollinator conservation. Future research should explore whether bee body-size is indeed relevant to flower petal size and foraging distance, and if we might use bee size to understand more about where bees are nesting so as to preserve and improve habitat spaces. We also plan to extend similar methods to explore pollination in other cropping systems in the effort to protect pollinator communities and improve pollination of crops on a larger scale.

## Acknowledgements

We thank all of our grower partners and our funding sources (USDA AFRI #2018-67011-28021, SARE Grant # SW 18-031 and the WOCS Project # 3018. We thank K. Sowers, R. Bomberger, W.S. Sheppard, W. Mattingly, M. Brousil, B. Tobey, M. Sherwood, R. Ryan, S. Ball, C. Loch, B. Box, and P.B. Ironhorse for their support and contributions to this project.

## Works cited

Adamidis, G. C., R. V. Cartar, A. P. Melathopoulos, S. F. Pernal, and S. E. Hoover. 2019. Pollinators enhance crop yield and shorten the growing season by modulating plant functional characteristics: A comparison of 23 canola varieties. Sci. Rep. 9: 1–12.

Bjerke, J., B. Caldbeck, D. Crowder, C. Dalley, D. Epstein, J. Gunter, C. Hiatt, S. Hoover, J. Knodel, B. Nelson, R. Olsson, T. Royer, D. Thorenson, T. Steeger, and R. Verhoek. 2019. Best Management Practices (BMPs) for Pollinator Protection in Canola Fields. Washington D.C.

Black, A. E., J. M. Scott, E. Strand, P. Morgan, C. Watson, and G. Wright. 1998. Biodiversity and Land-use History of the Palouse BioregionLJ: Pre-European to Present Biodiversity and Land-use History of the Palouse BioregionLJ: Pre-European to Present by, L. Use Hist. North Am.

Brown, J., and J. Davis. 2020. Brassica Breeding and Research Goals. http://www.cals.uidaho.edu/brassica/goals.asp.

Canada, C. C. of. 2017. History of Varietal Development. (https://www.canolacouncil.org/canola-encyclopedia/crop-development/history-of-varietal-development/#low-erucic-acid-rapeseed-varieties).

Cane, J. H., R. L. Minckley, L. J. Kervin, T. H. Roulston, and N. M. Williams. 2006. Complex responses within a desert bee guild (Hymenoptera: Apiformes) to urban habitat fragmentation. Ecol. Appl. 16: 632–44.

Carruthers, J. M., S. M. Cook, G. A. Wright, J. L. Osborne, S. J. Clark, J. L. Swain, and A. J. Haughton. 2017. Oilseed rape (Brassica napus) as a resource for farmland insect pollinators: quantifying floral traits in conventional varieties and breeding systems. GCB Bioenergy. 9: 1370–1379.

Chambó, E. D., N. T. E. de Oliveira, R. C. Garcia, M. C. C. Ruvolo-Takasusuki, and V. A. Arnaut de Toledo. 2017. Statistical modeling of insect behavioral response to changes in weather conditions in Brassica napus L. Arthropod. Plant. Interact. 11: 613–621.

Esser, A., and R. Hennings. 2012. Winter Canola Feasibility in Rotation with Winter Wheat. 1–4.

Farkas, Á., R. Molnár, T. Morschhauser, and I. Hahn. 2012. Variation in nectar volume and sugar concentration of allium ursinum L. Ssp. Ucrainicum in three habitats. Sci. World J. 2012: 7.

Finke, D. L. 2012. Contrasting the consumptive and non-consumptive cascading effects of natural enemies on vector-borne pathogens. Entomol. Exp. Appl. 144: 45–55.

Garnier, S. 2018. viridis: Default Color Maps from “matplotlib.”

Gathmann, A., and T. Tscharntke. 2002. Foraging ranges of solitary bees. J. Anim. Ecol. 71: 757–764.

Glozier, K. 2008. Protogol for Leaf Image Analysis. (https://ucanr.edu/sites/fruittree/files/49325.pdf).

Goodrich, B. 2019. Contracting for Pollination Services: Overview and Emerging Issues. Choices. 34: 1–13.

Goulet, H., and J. T. Huber. 1993. Hymenoptera of the world: An identification guide to families, Museum.

Goulson, D., E. Nicholls, C. Botías, and E. L. Rotheray. 2015. Bee declines driven by combined stress from parasites, pesticides, and lack of flowers. Science (80-.). 347: 1255957.1–9.

Greenleaf, S. S., N. M. Williams, R. Winfree, and C. Kremen. 2007. Bee foraging ranges and their relationship to body size. Oecologia. 153: 589–96.

Halinski, R., C. F. Dos Santos, T. G. Kaehler, and B. Blochtein. 2018. Influence of wild bee diversity on canola crop yields. Sociobiology. 65: 751–759.

Ibanez, S. 2012. Optimizing size thresholds in a plant-pollinator interaction web: Towards a mechanistic understanding of ecological networks. Oecologia. 170: 233–242.

Kahle, D., and H. Wickham. 2013. ggmap: Spatial Visualization with ggplot2. R J. 5: 144–161.

Klein, A.-M., B. E. Vaissière, J. H. Cane, I. Steffan-Dewenter, S. a Cunningham, C. Kremen, and T. Tscharntke. 2007. Importance of pollinators in changing landscapes for world crops. Proc. Biol. Sci. 274: 303–13.

Koenig, R., W. Hammac, and W. Pan. 2011. Canola growth, development, and fertility. Washingt. Washingt. State Univ. 1–6.

Michener, C. D. 2007. Bees of the World, 2nd ed. Johns Hopkins University Press, Baltimore.

Mohler, C. L. C., S. E. S. Johnson, and N. Resource. 2009. Crop Rotation on Organic Farms: A Planning Manual. NRAES, SARE, Ithaca.

Murcia, C. 1990. Effect of floral morphology and temperature on pollen receipt and removal in Ipomoea trichocarpa. Ecology.

O’Brien, C., and H. S. Arathi. 2018. Bee genera, diversity and abundance in genetically modified canola fields. GM Crop. Food. 9: 31–38.

Oksanen, J., F. G. Blanchet, M. Friendly, R. Kindt, P. Legendre, D. McGlinn, P. R. Minchin, R. B. O’Hara, G. Simpson, L.P. Solymos, M. Henry, H. Stevens, E. Szoecs, and H. Wagner. 2019. vegan: Community Ecology Package.

Olsson, R. L., K. Sowers, and Crowder. 2021. POLLINATORS IN CANOLA IN THE INLAND PACIFIC NORTHWEST Pollinators in Canola in the Inland Pacific Northwest. WSU PNW Publ. 1–13.

Pan, W. L., F. L. Young, T. M. Maaz, and D. R. Huggins. 2016. Canola integration into semi-arid wheat cropping systems of the inland Pacific Northwestern USA. Crop Pasture Sci. 67: 253–265.

Petanidou, T., A. S. Kallimanis, J. Tzanopoulos, S. P. Sgardelis, and J. D. Pantis. 2008. Long-term observation of a pollination network: Fluctuation in species and interactions, relative invariance of network structure and implications for estimates of specialization. Ecol. Lett. 11: 564–575.

Rasband, W. n.d. ImageJ.

Rhoades, P. R., T. Griswold, H. Ikerd, L. Waits, N. Bosque-Pérez, and S. Eigenbrode. 2017. The native bee fauna of the Palouse Prairie (Hymenoptera: Apoidea). J. Melittology. 83844: 1–20.

Team, R. C. 2019. R: A language and environment for statistical computing.

Torné-Noguera, A., A. Rodrigo, X. Arnan, S. Osorio, H. Barril-Graells, L. C. Da Rocha-Filho, and J. Bosch. 2014. Determinants of spatial distribution in a bee community: Nesting resources, flower resources, and body size. PLoS One. 9: 1–10.

Ullmann, K. S., M. H. Meisner, and N. M. Williams. 2016. Impact of tillage on the crop pollinating, ground-nesting bee, Peponapis pruinosa in California. Agric. Ecosyst. Environ. 232: 240–246.

USDA-NASS. 2019. Crop Production 2018 Summary.

Wickham, H. 2016. ggplot2: Elegant Graphics for Data Analysis.

Wiemer, a. P.a., N. Sérsic, S. Marino, a. O. Simões, and a. a. Cocucci. 2012. Functional morphology and wasp pollination of two South American asclepiads (Asclepiadoideae-Apocynaceae). Ann. Bot. 109: 77–93.

WOCS. 2017. Washington Oilseed Cropping Systems Project.

